# Gene silencing in *Cryptosporidium*: A rapid approach to identify novel targets for drug development

**DOI:** 10.1101/626317

**Authors:** A Castellanos-Gonzalez, G Martinez-Traverso, K Fishbeck, S Nava, AC White

## Abstract

**Background:** Cryptosporidiosis is a major cause of diarrheal disease. However, the only drug approved for cryptosporidiosis does not work well in high risk populations. Therefore, novel drugs are urgently needed. Then, the identification of novel is necessary to develop new therapies against this parasite. Recently, we have developed a rapid method to block gene expression in *Cryptosporidium* by using pre-assembled complexes of *Cryptosporidium* antisense RNA and human protein with slicer activity (Argonaute 2). We hypothesized that structural proteins, proteases, enzymes nucleotide synthesis and transcription factors are essential for parasite development, thus in this work we knock down expression of 4 selected genes: Actin, Apicomplexan DNA-binding protein (AP2), Rhomboid protein 1 (Rom 1) and nucleoside diphosphate kinase (NDK) and elucidated its role during invasion, proliferation and egress of *Cryptosporidium*.

**Methods:** We used protein transfection reagents (PTR) to introduce pre-assembled complexes of antisense RNA and human Argonaute 2 into *Cryptosporidium parvum* oocysts, the complexes blocked expression of Actin (Act), Transcription factor AP2 (AP2), nucleoside diphosphate kinase (DKN), and rhomboid protein 1 (Rom1). After gene silencing, we evaluated parasite reduction using *In vitro* models of excystation, invasion, proliferation and egress. We evaluated the potency of ellagic acid, a nucleoside diphosphate kinase inhibitor for anti-cryptosporidial activity using a model of *in vitro* infection with human HCT-8 cells.

**Results:** Silencing of Act, AP2, NDK and Rom1 reduce significantly invasion, proliferation and egress of *Cryptosporidium*. We showed that silencing of NDK markedly inhibited *Cryptosporidium* proliferation. This was confirmed by demonstration that ellagic acid reduced the number of parasites at micro molar concentrations (EC 50 =15-30 µM) without showing any toxic effect on human cells.

**Conclusions:** Overall the results confirmed the usefulness RNA silencing can be used to identify novel targets for drug development against *Cryptosporidium*. We identified ellagic acid (EA), a nucleoside diphosphate kinase inhibitor also blocks *Cryptosporidium* proliferation. Since EA is a dietary supplement approved for human use, then this compound should be studied as a potential treatment for cryptosporidiosis.

**Author summary:** The World Health Organization reports diarrhea kills around 760,000 children under five every year. *Cryptosporidium* infection is a leading cause of diarrhea morbidity and mortality. Current therapies to treat this infection are suboptimal, therefore novel treatments are urgently needed. We used genetic tools to identify novel targets for drug development, thus in this work we evaluated the role of 4 genes during *Cryptosporidium* infection. We demonstrated that silencing of nucleoside-diphosphate kinase (NDK) drastically reduced invasion, proliferation and egress of this parasite. To validate these finding we used the Ellagic acid (EA) an inhibitor of NDK to treat infected intestinal cells. Our results confirmed that the EA blocks parasite proliferation on infected cells. Interestingly we observed that the ellagic acid also has anti cryptosporidial activity by inducing apoptosis. Since EA is a dietary supplement already approved for human use, then this compound has potential to be used as a rapid alternative to treat Cryptosporidiosis.

## Introduction

*Cryptosporidium* is a leading cause of moderate-to-severe diarrhea in children under two years old and the leading pathogen associated with death in toddlers (ages 12 to 23 months^1^). Nitazoxanide is only one FDA approved medicine available for cryptosporidiosis, but it has limited efficacy in the population at the highest risk for poor outcomes. There is a strong consensus that better treatment options are urgently needed^2-4^. The limitation of tools to genetically manipulate gene expression in this parasite has been identified as a major hurdle for drug and vaccine development^2,4^. We developed a method to silence genes in this parasite by using preassembled complexes of *Cryptosporidium* single strand RNA and the human enzyme Argonaute 2 (hAgo2)^5^. We hypothesized that this method could be used to study key steps of infection and identify novel targets for drug and vaccine development. In the present work we used gene silencing to evaluate the role of Actin (Act), Rhomboid protein 1 (Rom1), transcription factor AP2 (AP2) and Nucleoside diphosphate kinase (NDK) 1 during *Cryptosporidium* infection.

## Methods

### Target selection for silencing experiments

Initially, we selected 100 genes for silencing experiments (Table S1), for these experiments mRNA sequences were obtained from CryptoDB (https://cryptodb.org/) and Gene Bank data bases (https://www.ncbi.nlm.nih.gov/nucleotide/). Selected genes codes for: structural proteins, transcription factors, kinases and proteases (Table S1). In these experiments we observed silencing ranging from 30-94%, however in this study we only analyzed genes that were silenced >75% (Table 1, figure 1).

**Table 1.** Gene silencing in *Cryptosporidium*. Selected targets (yellow column) were silenced with antisense ssRNA (gray column) and hAgo2 complexes. Silencing was evaluated by RT-PCR and Ct values were determined in treated with hAgo2/target ssRNA (red column) and hAgo2/unrelated ssRNA (green column). The silencing was calculated as fold change relative to control sample and expressed as percentage, SD=standard deviation (blue column).

| RT-PCR<br>amplified<br>Target | Antisense<br>ssRNA<br>21nt | Ct value<br>Ago/ssRNA | Ct value<br>Ago/unrelated<br>RNA | % silencing<br>± SD |
| --- | --- | --- | --- | --- |
| Actin | ssAct | 31.4 ± 0.4 | 28.3 ± 0.8 | 78 ± 3 |
| Ap2 | ssAp2 | 25.3 ± 0.3 | 21.3 ± 0.2 | 94 ± 0 |
| NDK | ssNDK | 27.3 ± 0.7 | 23.6 ± 0.6 | 93 ± 0 |
| Rom1 | ssRom1 | 46.5 ± 4.1 | 43.0 ± 2.7 | 96 ± 1 |

**Figure 1.**
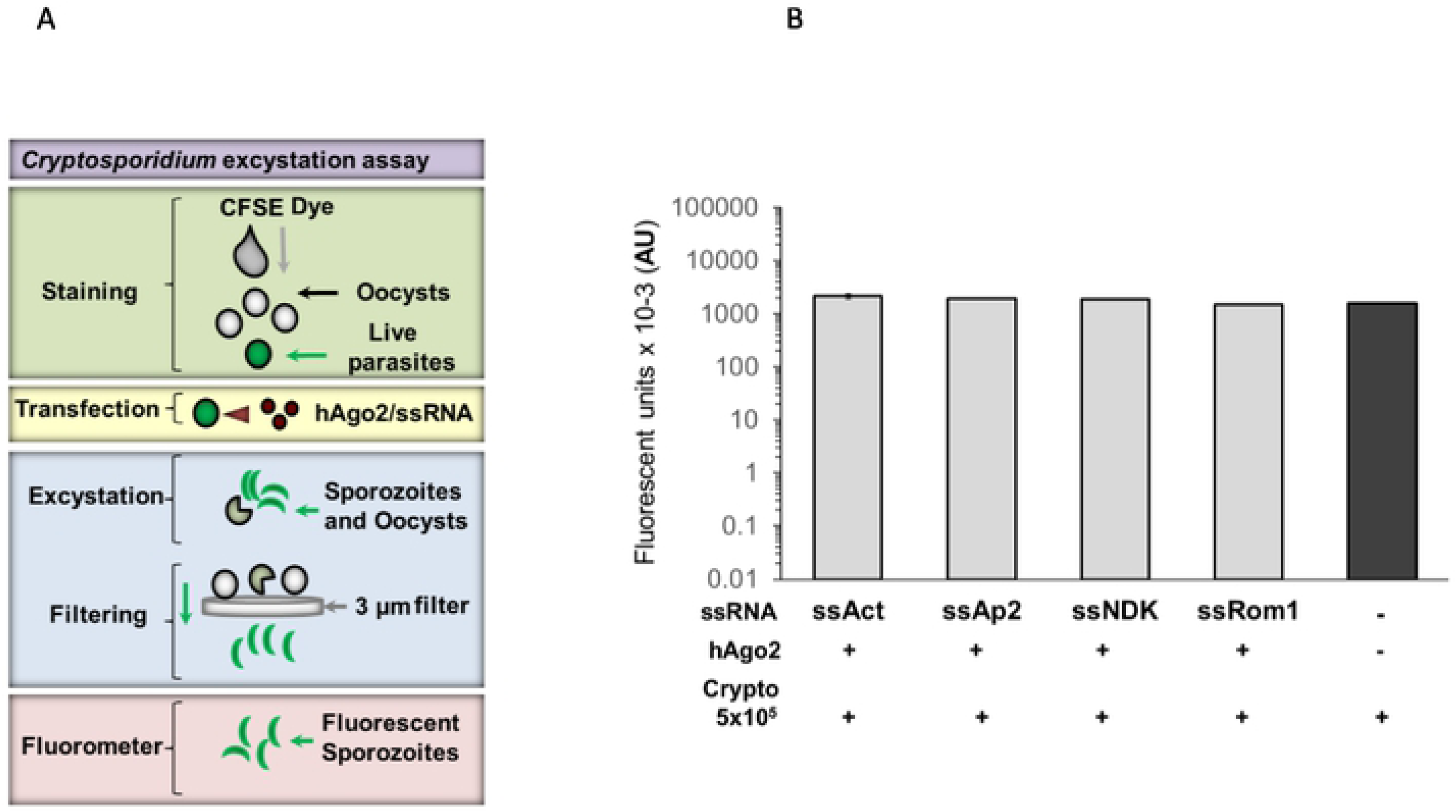
*Cryptosporidium* excystation assay. *Excystation model (A).* Parasites were stained with a vital dye (highlighted in green) and transfected with hAgo2/target ssRNA (highlighted in yellow), then excystation was induced and sporozoites were filtered (highlighted in blue). Sporozoites were quantified by fluorometry (highlighted in purple). *Effect of silencing on excystation (B). Cryptosporidium* oocysts were transfected with silencer complexes (+) or not (-), we induced excystation and compared fluorescence in treated (grey bars) and untreated oocysts (black bar). Experiments were conducted by triplicate. Non statistical difference was observed between control and treated samples.

### Antisense ssRNA design

Antisense single stranded RNA (ssRNA) used in silencing experiments was designed by using the computational software sFold 2.2 (http://sfold.wadsworth.org). We used as template the full sequences of mRNA targets (accession numbers in table S1), initially we generated all possible ssRNA antisense sequences of 21 nucleotides for each target, however we only selected optimal ssRNAs based in sFold ranking, these scores reflects parameters such as, local free energy and binding probability (C-G>40%). We synthetized selected ssRNAs from a commercial vendor (Integrated DNA Technologies, Coralville, IA), for silencing experiments the ssRNA was modified as follow: 21 nt ssRNA was capped with a phosphorylation at 5’ end and was modified with a deoxinucleotide (dTdT) tail at 3’ end (Table S2). Scrambled control ssRNA (Table S2) was designed using siRNA wizard software (Invivogen, San Diego CA).

### Gene silencing in C. parvum oocysts

For transfection experiments, we used oocysts (Iowa isolate) purchased from University of Arizona (Sterling Laboratories, Tucson, AZ). First the oocyst were prepared for transfection: the oocysts (1×10^6^ for each target) were transferred to 1.5 ml tubes, the samples were diluted with nuclease free water (Fisher Scientific, Hampton, NH) and then centrifuged for 10 minutes at a speed of 8,000 rpm (using microcentrifuge Eppendorf 5424). The supernatant was discarded and then the pellet was resuspended with 20 µl of nuclease-free water and then the tubes with the sample were placed at RT before adding transfection reagents. To assemble silencer complexes, first the ssRNAs were diluted with water at 100 nM, then samples were heated for 1 minute at 95°C and placed on ice. Complexes of ssRNA-hAgo2 were assembled in 1.5 ml microcentrifuge tubes, each tube contained 2.5 µl of diluted ssRNA 100 nM, 2.5 µl [62.5 ng/ul] of human Argonaute 2 (hAgo2) protein (Sino Biologicals, North Wales, PA) and 15 µl of Assembling Buffer [2 mM Mg(OAc)_2_, 150 mM KOHAc, 30 mM HEPES, 5 mM DTT, nuclease-free water]. The mixture was incubated for 1 hour at RT. After incubation, the complexes were encapsulated by adding 15 µl of protein transfection reagent (PTR) Pro-Ject™ (Thermo Scientific, Rockford, IL). The sample was mixed by pipetting and then incubated for 30 min, at RT. For transfection experiments the encapsulated complexes were added to oocysts and incubated at room temperature for 2 hrs. The slicer activity of hAgo2 was activated by incubating at 37 °C for 2 hrs. The reaction was stopped by adding 350 µl RLT lysis buffer (RNeasy kit, Qiagen, Hilden, Germany) and then samples were stored at −20°C for its posterior analysis by RT-PCR. For some experiments we used only PTR or PTR with unrelated ssRNAs, scrambled ssRNA (Table S2).

### RNA extraction and evaluation of silencing by RT-PCR

Prior to RNA isolation, samples (previously stored at −20°C) were thawed at 95°C for 2 minutes. Then, the total RNA was extracted from samples with the Qiagen’s RNeasy Plus Mini Kit (Qiagen, Valencia CA) following the instructions of the vendor. The RNA was eluted from purification columns with 100 µl of RNase-free water, then the concentration of eluted RNA was determined by spectrophotometry using a NanoDrop 100 Spectrophotometer (Thermo Fisher Scientific, Waltham MA). The silencing in transfected oocysts was analyzed by qRT-PCR using qScript™ One-Step SYBR® Green qRT-PCR Kit, Low ROX™ (Quanta BioSciences/VWR, Radnor, PA). For RT-PCR experiments, reactions were assembled as follow: 2 µl of purified RNA template [20 ng/µl], 5 µl of the One-Step SYBR Green Master Mix, 0.25 μl of each primer at a 10 μM concentration, 0.25 μl of the qScript One-Step reverse transcriptase, and 4.25 µl of nuclease-free water for a total of 10 μl of mix per sample. The RT-PCR mixture (total volume 12 µl) was transferred to 96-well Reaction Plates (0.1 mL) (Applied Biosystems, Foster City, CA) and then RT-PCR amplification was conducted on a 7500 Fast Real-Time PCR System (Applied Biosystems, Foster City, CA) with the following cycling conditions: 50°C for 15 minutes, 95°C for 5 minutes, then 50 cycles of 95°C for 15 seconds and 63°C for 1 minute, followed by a melting point analysis (95°C for 15 seconds, 60°C for 1 minute, 95°C for 15 seconds and 60°C for 15 seconds). Before fold change analysis all the target Ct values for each silenced target were normalized against the *Cryptosporidium* GAPDH. To calculate fold changes between control samples and silenced samples, we used the ΔΔCt method. Results are shown by the average for each target with the respective standard deviation. List of primers used for RT-PCR is indicated in table S3.

### Oocyst excystation assays

Excystation of transfected oocysts was induced with acidic water and taurocholic acid as described before. Briefly, *Cryptosporidium* oocysts were pelleted by centrifugation (500 g), supernatant was discarded and then parasites were resuspended in 25 µl acidic water (pH 2.5), and incubated for 10 minutes on ice. Then, excystation media 250 µl (RPMI-1640 media, 1X antibiotic/antimycotic solution and 0.8% taurocholic acid sodium salt hydrate] was added. The sample was incubated for 1 hour at 37°C. After excystation we evaluate excystation rate, then the sporozoites were stained with the vital dye carboxyfluorescein succinimidyl ester (CFSE) (CellTraceTM, Thermo Fischer Scientific, Waltham, MA) by adding 2 µM of CFSE and incubating in the dark at 37°C for 15. After staining, the sporozoites were separated from unhatched oocysts by filtration using 3.0 µm nitrocellulose membranes (Merck Millipore Ltd., County Cork, Ireland). Fluorescence of filtered samples was evaluated, for these experiments 200 µl of each filtered sample was transferred to a 96-well plates (Costar, Corning, NY) and then fluorescence was measured with a microplate reader, (FLUOstar Omega, Ortenberg, Germany).

### HCT8 cell culture

For infection experiments we used ileocecal cells (HCT-8 cells, ATCC, Manassas, VA). For these experiments cells were thawed at 37°C and then resuspended with 500µl of RPMI-1640 media (Gibco/Thermo Fisher Scientific, Waltham, MA) supplemented with 10% fetal bovine serum (FBS) (Stemcell Technologies, Vancouver, Canada) and 1X antibiotic/antimycotic solution (Gibco/Thermo Fisher Scientific, Waltham, MA) and plated in a 24-well plate (Costar, Corning, NY). Cells were incubated at 37°C overnight.

### In vitro Invasion assay

For invasion assay HCT8 cells were cultured as described. Before the infection, transfected parasites were stained and excysted as described. After filtering, approximately 5×10^5^ (suspended in 250 µl) were added to HCT-8 cells for 1 hr at 37°C. To quantify sporozoites that did not invade cells, 250 µl of the supernatant was collected after incubation and fixed with 50 µl of 4.2%paraformaldehyde solution (Cytofix/Cytoperm, BD BioSciences, San Jose CA). To quantify sporozoites adhered to the cells, the monolayers were trypsinized by adding 150 µl of 0.25% trypsin-EDTA (Gibco/Thermo Fisher Scientific, Waltham, MA), and incubating at 37°C for 15 minutes. Then, the trypsin was inactivated by adding 500 µl of RPMI media supplemented with 10% FBS. Samples were transferred to 1.5 ml tubes and then centrifuged 10 min at 500 g, supernatant was removed, and pellet was resuspended in 50 µl of Cytofix solution. Fixed sample was resuspended in 200 µl of 1X PBS (Fisher Scientific, Fair Lawn, NJ) and was filtered using a 5 ml Falcon polystyrene round-bottom tube with a cell-strainer cap with a 35 µm nylon mesh (Corning Inc., Corning, NY). Filtered samples were analyzed by flow cytometry using a SE500 Flow Cytometer (Stratedigm, San Jose CA). To define sporozoites populations on infected cells, we analyzed filtered sporozoites stained with CFSE but without HCT-8 cells (Fig 1A).

### In vitro Proliferation assay

The vital dye, CFSE (Thermo Fisher Scientific, Waltham, MA) was used to track proliferation tracker of intracellular stages of *C. parvum*. CFSE is activated by viable cells. However, with each cycle of cell division, the fluorescent intensity decreases. Thus, proliferation was evaluated by measuring the reduction of fluorescence intensity by flow cytometry after 16 hours (Fig 3A and 3B). For these experiments the parasites were silenced (or not) and excysted as described before. After excystation, sporozoites were used to infect HCT-8 cells cultured as previously described. Basal infection was allowed for 2 hours at 37°C. The monolayer was then washed and media replaced with 250 µl of fresh RPMI-1640 with 10% FBS and 1X antibiotic/antimycotic solution. Infected cells were incubated at 37°C for 16 hours (before parasite egress at 19-24 hrs). After incubation, cells were harvested by trypsinization and then washed, fixed, resuspended in 1X PBS and analyzed by flow cytometry.

**Figure 2.**
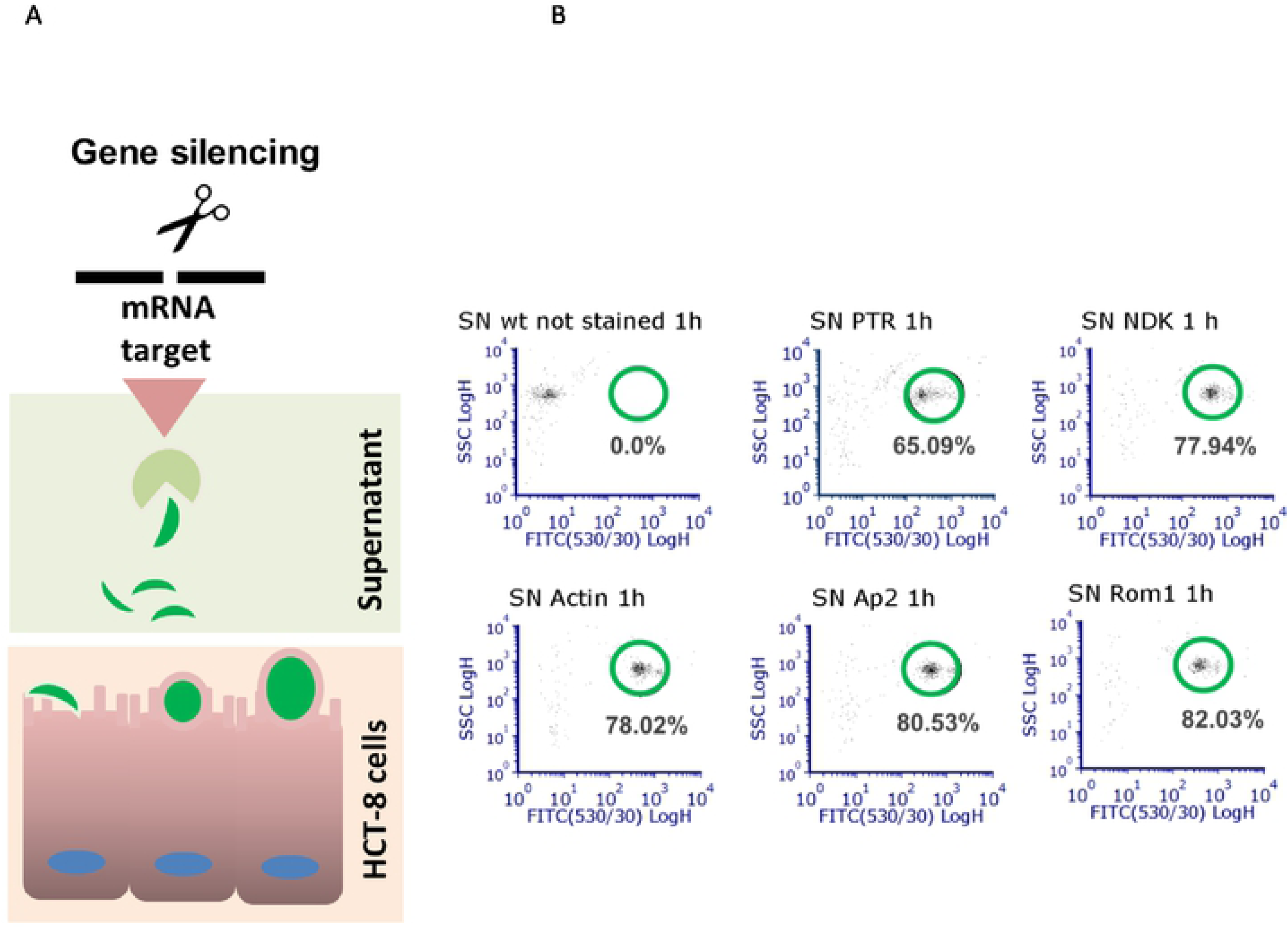
*Cryptosporidium* invasion assay. *Invasion model (A).*Transfected parasites were used to infect HCT-8 cells (infected cells shown in red square), after 1 hr of infection supernatant (square green) was collected and % of sporozoites was evaluated by flow cytometry. *Effect of silencing on parasite entrance by Flow cytometry (B).* Silencing was induced or not (PTR) and then sporozoites were used to infect cells, parasites that did not invade cells were evaluated in supernatants (SN). wt= unstained wild typte, PTR= parasites treated only with protein transfection reagent.

**Figure 3.**
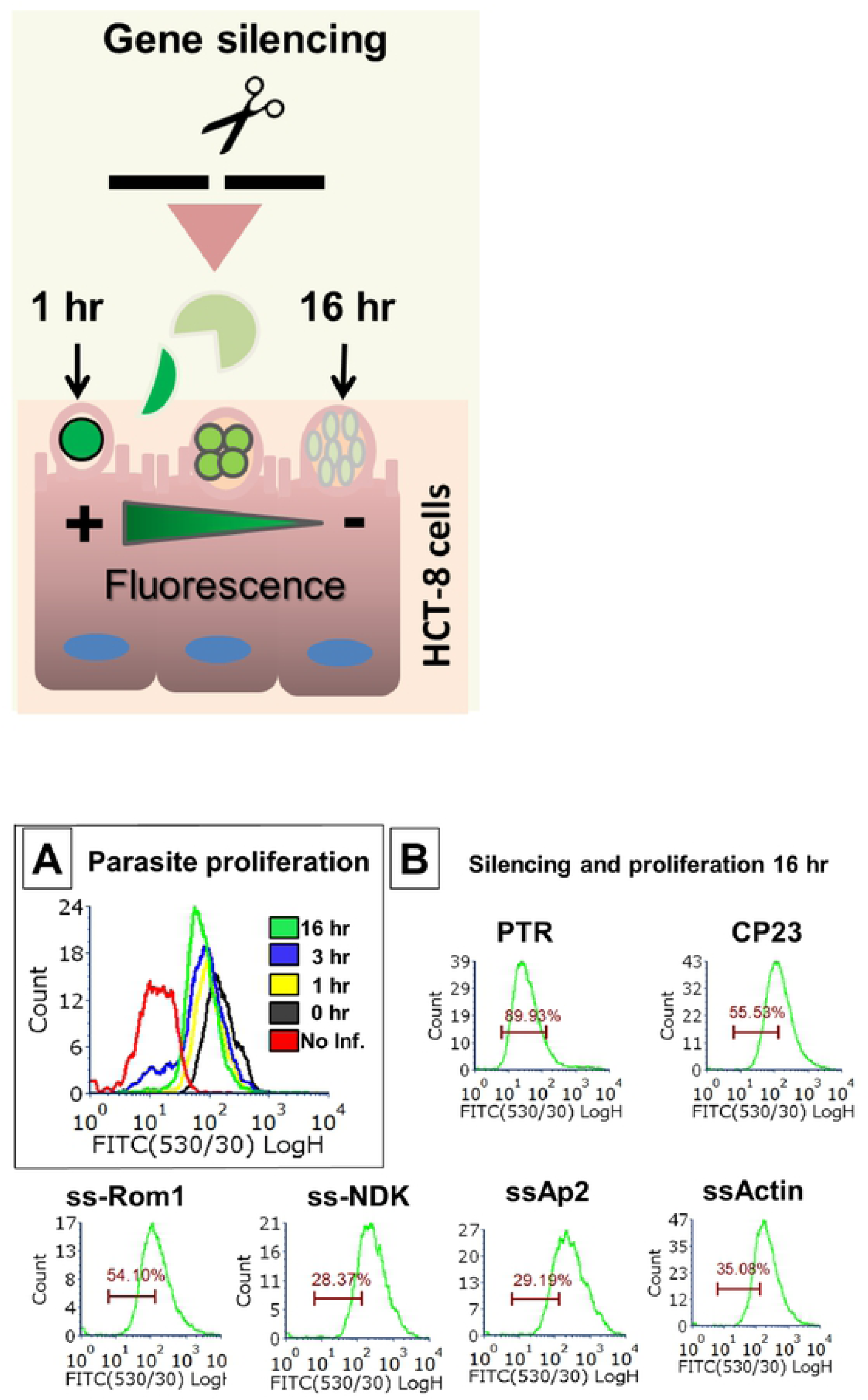
*Cryptosporidium* proliferation assay. Stained parasites (sowed in green) were transfected and then used to infect HCT-8 cells, division of WT parasites within infected cells was analyzed by flow cytometry 0-16 hrs tracking reduction of fluorescent signal (A). Division of transfected parasites was analyzed at 16 hrs and compared with untreated parasites (PTR) or parasites treated with unrelated target Cp23 (B).

### Merozoite egress assay

We evaluated the effect of silencing using an egress model by measuring the number of merozoites in supernatants collected between 16-19 hrs post-infection. For these experiments we induced the silencing after infection of the HCT-8 cells. After sporozoite infection of HCT-8 cells, media was removed and cells were transfected with ssRNA/hAgo2 complexes and incubated at 37°C for 16 or 19 hours. RPMI media was removed at 16 hours, replaced with 250 µl of fresh RPMI media, and incubated for 3 hours more to complete the 19 hours incubation period. After incubations, the supernatant was collected, filtered and fixed as before. After the supernatant was removed, the remaining cell monolayer was trypsinized, washed, fixed and filtered. The supernatant and the cells monolayer were separately analyzed by flow cytometry.

### Anticryptosporidial activity of ellagic acid on infected cells

To evaluate anticryptosporidial effect of ellagic acid HCT-8 cells were infected with CFSE-labeled sporozoites. After 1 hour of infection, media was replaced with 250 µl of serum-free RPMI media (free serum) containing varying concentrations of ellagic acid [0, 3, 30, 300 nM and 3µM] and incubated the sample for 16 hrs. After the incubation, the monolayers in the plate were washed with PBS. The cells were trypsinized, fixed and analyzed by flow cytometry as described. For RT-PCR experiments, monolayers were washed with PBS. After removing the supernatant, 350 µl of RLT buffer was added and RNA extracted with RNAeasy kit

## Results

### Silencing of Actin, Rom1, NDK, and TB2

Four (21nt) ssRNA antisense sequences each were synthesized complementary to mRNA of Actin, Rom1, DKN, and TB2 (Table S2). Complexes for all 4 targeted genes led to >75% decreased expression when compared to controls (unrelated ssRNA, scramble ssRNA or untreated parasites, (Table 1). ssRNA-Ago did not affect the expression of non-targeted GAPDH mRNA, ribosomal r18s, or parasite viability (Fig. S2).

### Silenced targets are not involved on excystation of Cryptosporidium parasites

We evaluated the role of silenced genes during excystation of *Cryptosporidium* sporozoites measuring the excystation rate by fluorescence (Fig 1A). Silencing did not produce a significant reduction on excystation rate for any of the targets (Fig. 1B).

### Gene silencing of selected genes blocks parasite entry

We evaluated gene silencing by flow cytometry using an invasion model (Fig 2A), for these experiments we measured the proportion of CSFE-labeled parasites that failed to invade HCT-8 cells (Fig 2A). First, we defined sporozoites population by flow cytometry (Fig S1A). For the invasion model, we transfected parasites and then evaluated the number of sporozoites in the supernatant s. The results indicates that control group transfected only with PTR has ∼65% of stained cells (merozoites), in contrast silencing of Rom1 significantly increased the number of gated cells, meaning that sporozoites did not invaded the host cells (Fig 2B). Silencing of NDK, Ap2 and Actin also showed a partially effect on sporozoite invasion (Fig 2B). We did not observed differences between untreated parasites and transfected parasites with PTR (data not shown).

### NDK inhibition reduces parasite proliferation

To evaluate the effects of gene silencing on parasite division we used a proliferation model (Fig 3), parasites were labeled with CFSE and collected at 16 hours post-infection prior to the time of egress (Fig 3). For controls, cell proliferation led to decreased CFSE signal, such that 89% of cells had deceased signal by 16 hours (Fig 3A). By contrast, after silencing of NKD, Ap2 or Actin, only 28-35% of cells had decreased CFSE signal (Figure 3B). Silencing of CP23 and Rom1 had intermediate values, suggesting partial inhibition of proliferation (54-55%). In additional studies we tested the effect of EA acid on sporozoites viability, we did not observe killing effect of EA on sporozoites (Fig S1B).

### Silencing of Rom1 and AP2 reduced parasite egress

To test the effects of silencing on egress, we transfected intracellular parasites on infected HCT-8 cells. Transfected complexes did not affect the viability of HCT-8 cells (Fig S2), however we observed a reduction on expression in all tested targets (Fig S3). After confirmation of silencing, fresh media was added to collect merozoites released between 16-19 hrs (Fig S4). To evaluate the egress, we conducted qRT-PCR to quantify the relative number of merozoites in supernatants of treated samples and untreated samples (Fig 4B). There was a significant reduction in the number of merozoites observed in the supernatant of silenced samples (Fig 4B). Therefore, these results indicated that silencing of DKN and Actin reduced both parasite proliferation and egress. By contrast, silencing of Ap2 and Rom1 markedly reduced egress out of proportion to the effects on proliferation.

**Figure 4.**
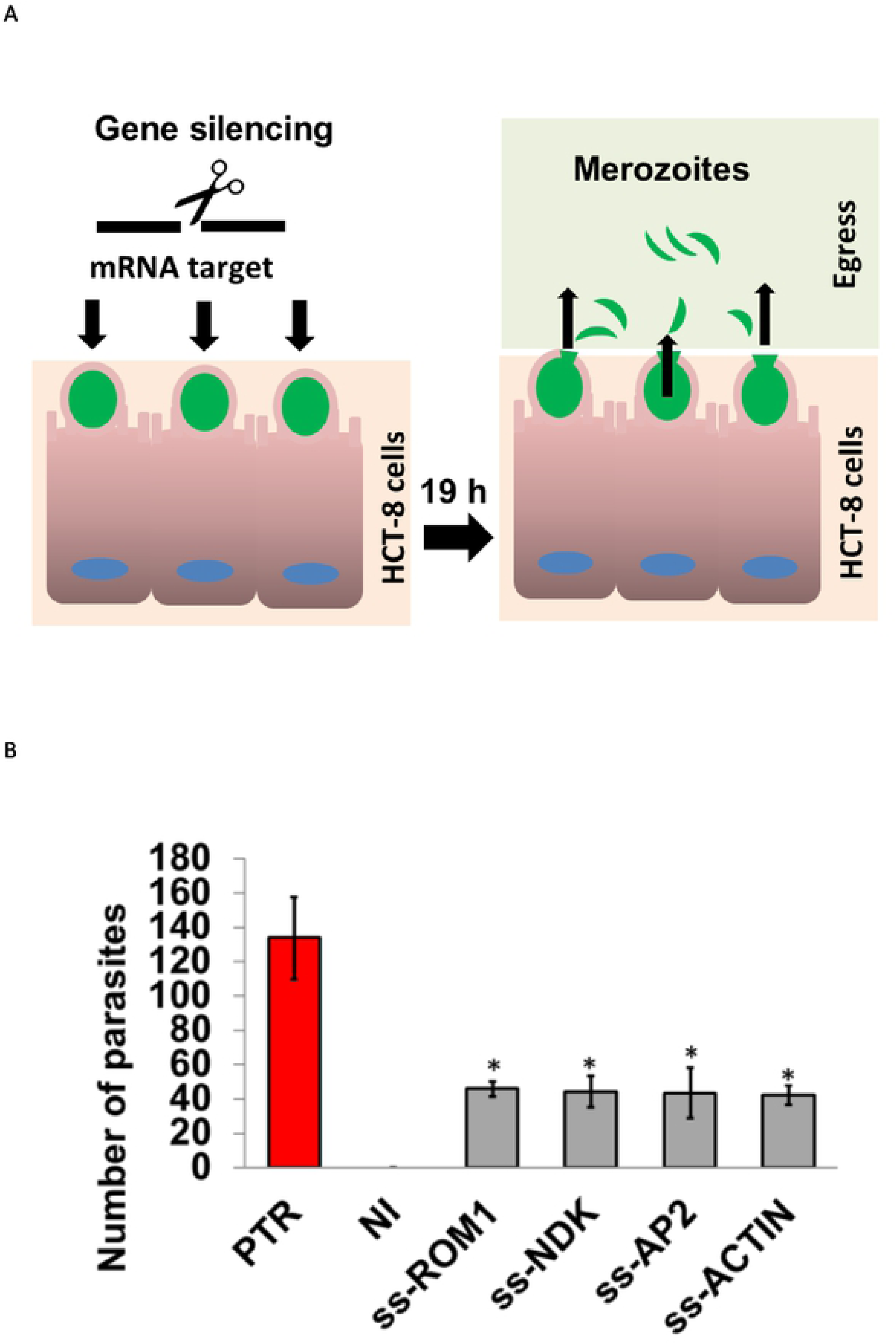
*Cryptosporidium* egress assay. Egress model (A). Parasites were stained (green) and used to infect HCT-8 cells. We induce silencing on parasites within infected cells. Then, supernatant from 16-19 hrs was collected and evaluated by flow cytometry (B). Total numbers of parasites (in 200 µl) were determined by qRT-PCR in treated samples (grey bars) and control sample treated only with PTR (red bar) and non-infected cells (NI). The experiments were conducted in triplicates, SD and p values (*= p≤ 0.05) are shown.

### Ellagic acid treatment blocks parasite proliferation

Since NDK silencing showed the highest level of reduction in proliferation, we evaluated the anticryptosporidial activity of the NDK inhibitor Ellagic acid (EA) in the HCT-8 infection model. The results showed that EA inhibited parasite proliferation at micromolar concentrations (Fig 5), with an EC50 of within 15-30 µM. In contrast, our results showed that EA is not cytotoxic at these concentrations (Supplementary figure S5). In order to evaluate the anticryptosporidial mechanism, we evaluated the expression of proliferation and apoptosis markers in the parasites. Our results showed a significant down regulation of separine and meta-caspases (Fig 5B-C), which suggest a parasitostatic and parasiticidal effect by blocking proliferation and inducing apoptosis.

**Figure 5.**
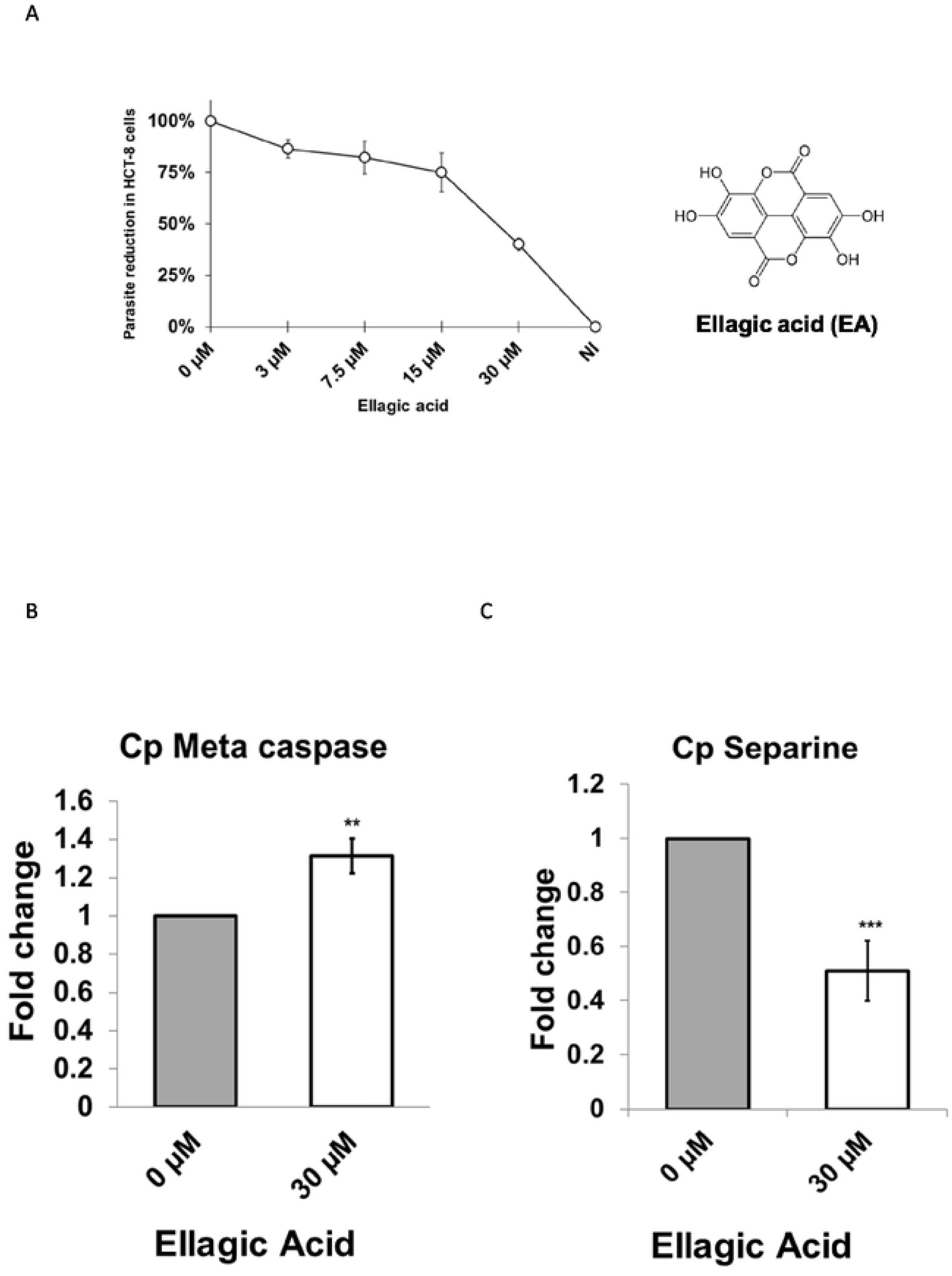
Anticryptosporidial activity of Ellagic acid (EA). Reduction of parasites on infected cells by EA. We infected human intestinal cells (HCT-8) with *Cryptosporidium*, after the infection cells were treated with different amounts of EA and then we evaluated the parasite burden by qRT-PCR (A). Activation of apoptosis by EA. We evaluated expression of apoptosis marker metacaspase in treated (with bar) and untreated (grey bar) cells (B). Inhibition of proliferation by EA. Expression of proliferation marker separine was evaluated by qRT-PCR on infected cells treated (with bar) and untreated (grey bar) with EA (C). RT-PCR experiments were conducted by triplicate, and standard deviation is indicated in each bar.

## Discussion

*Cryptosporidium* lacks the machinery involved with mammalian gene silencing^6^. In previous studies, we have demonstrated the feasibility to silence *Cryptosporidium* genes by transfecting oocysts with human Argonaute (with slicer activity) loaded with ssRNA^5^. These initial experiments confirmed reduction at protein levels and pointed out the usefulness of the method to evaluate parasite invasion. Since the silencing is maintained up to 24 hrs., then we hypothesized that this method could be used to evaluate other key biological processes during the asexual cycle of parasites maintained in HCT-8 cells (e.g. excystation, proliferation and egress). Thus the overall goal of this study was to use the silencing method to identify targets that are critical for different steps in the parasite life-cycle. Our first step in this work was to identify druggable candidates for gene silencing. We used transcriptional data to prioritize genes highly expressed during invasion, proliferation and egress stages. Also we prioritized genes with low homology with host molecules, but highly conserved between *Cryptosporidium* species. After initial analysis, we identified 100 potential candidates (Fig S1). We developed antisense ssRNA sequences to silence selected genes. In these experiments silencing rates were among 30-94% (Data not shown). Since partial silencing (30-75%) may not be optimal for phenotypic studies, then in this work we only used antisense ssRNA that induces >75%. Selected genes for silencing included: 1) Actin, an structural essential for *Cryptosporidium* motility^7^, 2) NDK an essential gene for synthesis of nucleotides ^8^ 3) Rom1, a protease involved on invasion and egress in other apicomplexan^9^ and 4) Ap2, which is a transcription factor involved in proliferation^7^. First, we evaluated the effect of silencing during parasite excystation. The excystation assays showed that none of the silenced genes had effect on this process suggesting that these proteins are not essential for excystation. This result was expected since transcriptomic data has showed that the majority of genes (∼85%) in *Cryptosporidium* are expressed after excystation process^10^. The invasion assay indicated that silencing of Rom1 blocks parasite entry. This proteolytic enzyme has previously been implicated in parasite invasion^11^. Orthologue rhomboid protease in *Toxoplasma* cleaves cell surface adhesins, and have been demonstrated that this protein is essential for invasion^11^. In *Plasmodium* PfROM1 and PfROM4 helped in merozoite invasion by catalyzing the intramembrane cleavage of the merozoite adhesin AMA1^11^. Actin silencing also showed inhibited Invasion. Apicomplexan parasites actively invade host cells using a mechanism predicted to be powered by a parasite actin-dependent myosin motor. Actin in invasion was first suggested by studies demonstrating the ability of the actin polymerization inhibitor cytochalasin D (CytD) to block invasion^12^. The proliferation assay showed that NDK, Ap2, Actin but not Rom1 reduced parasite proliferation. Nucleoside diphosphate kinases (NDK) are enzymes required for the synthesis of nucleoside triphosphates (NTP) other than ATP. They provide NTPs for nucleic acid synthesis, CTP for lipid synthesis, UTP for polysaccharide synthesis and GTP for protein elongation, signal transduction and microtubule polymerization. Not surprisingly, NDK is essential for intracellular parasite proliferation. Actin proteins are related with cytoskeleton motility during cell division.

*Cryptosporidium* transcriptomic analysis demonstrated that actin is highly expressed between 12-48 hrs after the infection, during this time the parasite is actively dividing passing from a single cell (trophozoite) to 8 cells (meronts II) in 24 hrs. We showed that silencing of AP2 transcription factor also affected proliferation (Fig 3B). AP2 proteins are transcription factors which harbor a plant-like DNA-binding domain. Five AP2 proteins have been identified as key stage-specific regulators in *Plasmodium*, thus AP2 proteins have been implicated in *P. falciparum* var gene regulation by binding the SPE2 DNA motif and acting as a DNA-tethering protein involved in formation and maintenance of heterochromatin^13^. The role of AP2 proteins in gene regulation has also been investigated to a lesser degree in *T. gondii*, where several AP2 proteins have been implicated in regulating progression through the cell cycle ^14^ as well as crucial virulence factors^15^. Other studies have implicated AP2s in regulating a developmental transition^16^. Radke et al. ^17^ recently characterized a *T. gondii* AP2 and showed that this molecule acts as a repressor of bradyzoite development. Our results showed that egress assay was partially affected by parasite proliferation (Fig 4B). Silencing of NDK, AP2 and Actin blocked proliferation leading to a reduction in the number of merozoites (Fig 3B). In contrast, silencing with Rom 1 had an even great effect on egress. Since that protein only had moderate effect in proliferation (Fig 3B), it is likely Rom1 is implicated in parasite egress through a proteolytic mechanism. In *Pla*smodium, release of merozoites from schizonts resulted in the movement of *Plasmodium* ROM1 from the lateral asymmetric localization to the merozoite apical pole and the posterior pole ^18^.

Overall our in vitro studies confirmed that silenced genes blocks proliferation and egress in *Cryptosporidium* parasites, therefore we hypothesized that chemical inhibitors against these enzymes should arrest *Cryptosporidium* proliferation on infected cells. Since silencing of NDK showed effect during invasion, proliferation and egress then was selected for further studies. The inhibition of NDK activity by EA have been demonstrated ^19,20^. Thus here we tested EA on the *Cryptosporidium* infection model, we observed anticryptosporidial activity of this compound at micromolar concentrations (Fig 5A). The EA is a natural compound found in strawberries and other fruits, thus this compound have been used as dietary complement to treat several diseases^21,22^, interesting recently this compound showed its antimicrobial activity in the gastrointestinal pathogen *Helicobacter pylori* ^23^. In order to investigate the mechanism of anticryptosporidial activity we evaluated expression proliferation (separine) and apoptosis markers (metacaspase) in the parasite. We observed a down regulation in separine expression (also known as separase) which is implicated chromatin regulation during meiosis and mitosis processes^24^, this finding suggest that EA may be blocking proliferation through NDK inhibition as observed in silencing experiments, however also we observed an up regulation of metacaspase which suggest that other mechanisms may be involved in parasite killing. EA has showed multiple benefits to human health trough enhancement of immune system or epithelial barrier ^25-27^, thus we speculate that EA has a dual effect on *Cryptosporidium* by reducing the infection trough the activation of host pathways (e.g. defensin secretion) and affecting essential enzymes on parasites. Interesting the micromolar concentrations tested here are under the biological concentration of EA acid commonly used in humans ^28^, therefore future studies will be focused to characterize the effect of EA and metabolites on intestinal cells of infected mice. If these studies confirm the anticryptosporidial activity and activation of host response then we anticipate that this compound could be implemented in a very near future in the treatment in humans infected with *Cryptosporidium*.

## ACKNOWLEDGMENTS

AC was supported by the Bill & Melinda Gates Foundation grant: OPP1161026 and by grant: 5R21AI12627502 from the National Institute of Allergy and Infectious Diseases, National Institutes of Health.

## Supporting Information Legends

### Table and Figures Legends (supplemental data)

Table S1. Accession numbers of *Cryptosporidium* sequences used for antisense ssRNA design.

Table S2. *Cryptosporidium* antisense ssRNA sequences used in this study.

Table S3. Target genes and primer sequences used for RT-PCR analysis.

Figure S1. Sporozoites of *Cryptosporidium* by Flow cytometry (A). Cryptosporidium oocysts were stained with CFSE and then excystation was induced. Parasites were filtered and then sample was analyzed by flow cytometry. Effect of EA on oocyst and excystation by flow cytometry (B), we treated stained oocyst with EA and then excystation was induced.

Figure S2. Expression of human 18s gene in HCT-8-cells after transfection. HCT-8 cells were infected with *Cryptosporidium* sporozoites and then transfected with ssRNA. For these experiments, after 2 hours of infection the supernatant was removed and new culture media was added. Next, silencer complexes were added to the infected cells and incubated overnight. To evaluate cytotoxic effects of ssRNA or HCT-8 we quantified 18s rRNA, if cells die this should be reflected as reduction of this marker. Figure shows no differences between wild type (WT), protein transfection reagent alone (PTR), and tested ssRNAs ssAct, ssAp2, ssNDK, ssRom1 (encapsulated in PTR).

Figure S3. Gene silencing after infection. In this experiment we tested the silencing on intracellular forms (merozoites) within HCT-8 cells. For these experiments we infected and treated cells as described above and we evaluated the silencing by qRT-PCR in total RNA from infected cells transfected with silencer complexes. The figure shows PCR CT values from duplicate experiments of transfected cells with silencer complexes (gray bar) and cells transfected only with PTR. All cells treated with ssRNA showed a delay in the amplification cycle when compared with controls. All samples were normalized against GAPDH. Standard deviation of PCR triplicates is shown in each bar.

Figure S4. *Cryptosporidium* merozoites collected at 16 hrs of infection. *Cryptosporidium* merozoites by flow cytometry. HCT-8 cells were infected with labeled sporozoites. After infection, silencing was induced, then at 16 hrs of infection (before egress) supernatant was removed and fresh media was added. Supernatant was collected again a 19 hrs and analyzed by flow cytometry, then % of merozoites were evaluated in samples treated only with PTR (left), merozoites are not observed in supernatants of non-infected samples (right).

Figure S5. Ellagic acid and viability on HCT-8 cells. Effect of EA on non-infected (NI) HCT-8 cells. HCT-8 cells were treated with EA (or not) and then viability was evaluated measuring the activation of the vital dye CFSE. The figure shows the % of cells positives for CFSE.

